# Afucosylated *Plasmodium falciparum*-specific IgG is induced by infection but not by subunit vaccination

**DOI:** 10.1101/2021.04.23.441082

**Authors:** Mads Delbo Larsen, Mary Lopez-Perez, Emmanuel Kakra Dickson, Paulina Ampomah, Nicaise Tuikue Ndam, Jan Nouta, Carolien A M Koeleman, Agnes L Hipgrave Ederveen, Benjamin Mordmüller, Ali Salanti, Morten Agertoug Nielsen, Achille Massougbodji, C. Ellen van der Schoot, Michael F. Ofori, Manfred Wuhrer, Lars Hviid, Gestur Vidarsson

## Abstract

IgG specific for members of the *Plasmodium falciparum* erythrocyte membrane protein 1(PfEMP1) family, which mediates receptor- and tissue-specific sequestration of infected erythrocytes (IEs), is a central component of naturally acquired malaria immunity. PfEMP1-specific IgG is thought to protect via inhibition of IE sequestration, and through IgG-Fc Receptor (FcγR) mediated phagocytosis and killing of antibody-opsonized IEs. The affinity of afucosylated IgG to FcγRIIIa is elevated up to 40-fold compared to fucosylated IgG, resulting in enhanced antibody-dependent cellular cytotoxicity. Most IgG in plasma is fully fucosylated, but afucosylated IgG is elicited in response to enveloped viruses and to paternal alloantigens during pregnancy. Here we show that naturally acquired PfEMP1-specific IgG is likewise markedly afucosylated in a stable and exposure-dependent manner, and efficiently induces FcγRIIIa-dependent natural killer (NK) cell degranulation. In contrast, immunization with a soluble subunit vaccine based on VAR2CSA-type PfEMP1 resulted in fully fucosylated specific IgG. These results have implications for understanding natural and vaccine-induced antibody-mediated protective immunity to malaria.

**Summary:** Afucosylated IgG has enhanced Fc-receptor affinity and functionality, and is formed specifically against membrane proteins of enveloped viruses. We show that this also applies to *Plasmodium falciparum* erythrocyte membrane-specific IgG induced by natural infection, but not by soluble PfEMP1 vaccination.

## Introduction

The most severe form of malaria is caused by the protozoan parasite *Plasmodium falciparum*. The disease is currently estimated to cost around 400,000 lives a year, mostly of young children and pregnant women in sub-Saharan Africa. In addition, nearly 900,000 babies are born with a low birth weight as a consequence of placental malaria (PM) (World Health Organization, 2020). The particular virulence of *P. falciparum* is related to the efficient adhesion of the infected erythrocytes (IEs) to host receptors in the vasculature, such as endothelial protein C receptor, intercellular adhesion molecule 1, and oncofetal chondroitin sulfate A (Bengtsson et al., 2013; Fried and Duffy, 1996; Lennartz et al., 2017; Turner et al., 2013), mediated by members of the protein family *P. falciparum* erythrocyte membrane protein 1 (PfEMP1), embedded in the membrane of IE (Hviid and Jensen, 2015). The sequestration of IEs can cause tissue-specific circulatory compromise and inflammation, which in turn can lead to severe and life-threatening complications such as cerebral malaria (CM) and PM (Jensen et al., 2020; Rogerson et al., 2007). Severe malaria in children has repeatedly been shown to be associated with parasites expressing particular subsets of PfEMP1, such as Group A and B/A (Jensen et al., 2004; Turner et al., 2013), whereas PM is strongly associated with parasites expressing VAR2CSA-type PfEMP1 (Salanti et al., 2004; Tuikue Ndam et al., 2005).

Acquired protective immunity to *P. falciparum* malaria is mainly mediated by IgG with specificity for antigens expressed by the asexual blood-stage parasites (Cohen et al., 1961). PfEMP1 is a key target (Hviid and Jensen, 2015), although antibodies to other blood-stage antigens, such as the merozoite-specific antigens glutamate-rich protein (GLURP), merozoite surface protein 1 and reticulocyte binding protein homolog 5, also contribute to naturally acquired protection (Conway et al., 2000; Douglas et al., 2011; Kana et al., 2017). Importantly, the selective protection from severe malaria that develops early in childhood, is related to acquisition of IgG specific for Group A and B/A PfEMP1 variants (Bull et al., 2000; Cham et al., 2010; Jensen et al., 2004). As a result, life-threatening complications are rare in teenagers and beyond in *P. falciparum* endemic regions. PM, which is caused by selective accumulation of VAR2CSA-positive IEs in the placenta from early in pregnancy (Ofori et al., 2018; Schmiegelow et al., 2017), constitutes an important exception to this rule. Only VAR2CSA mediates adhesion to placenta-specific chondroitin sulfate (Duffy et al., 2006; Salanti et al., 2004; Viebig et al., 2005). Because of this, and because antibodies specific for non-pregnancy-related types of PfEMP1 do not cross-react with VAR2CSA (Barfod et al., 2010; Salanti et al., 2004; Tuikue Ndam et al., 2006), primigravid women are immunologically naïve to VAR2CSA and therefore highly susceptible to PM, despite general protective immunity acquired during childhood. However, substantial IgG-mediated protection against PM is acquired in a parity-dependent manner, and PM is therefore mainly a problem in the first pregnancy (Fried and Duffy, 1996; Fried et al., 1998; Ricke et al., 2000; Salanti et al., 2004; Staalsoe et al., 2004).

Acquired immunity mediated by PfEMP1-specific IgG is generally thought to rely on their ability to interfere directly with IE sequestration (i.e., neutralizing, adhesion-inhibitory antibodies). However, antibody-mediated opsonization of IEs is a likely additional effector function of these antibodies, since the antibody response to most *P. falciparum* asexual blood-stage antigens (including PfEMP1) is completely dominated by the cytophilic subclasses IgG1 and (to a lesser extent) IgG3 (Megnekou et al., 2005; Piper et al., 1999). Nevertheless, the relative importance of neutralization and opsonization remains largely unexplored. Complement-mediated destruction of IgG-coated IEs does not seem important (Larsen et al., 2019), suggesting that IgG opsonization of IEs by IgG functions mainly through IgG-Fc receptor (FcγR)-dependent phagocytosis and antibody-dependent cellular cytotoxicity (ADCC) (Arora et al., 2018; Ataide et al., 2011; Marsh et al., 1989). The latter involves FcγRIIIa (Ravetch and Perussia, 1989; Scallon et al., 1989). Binding of IgG to FcγRIIIa critically depends on the composition of a highly conserved N-linked glycan at position 297 in the Fc region (Vidarsson et al., 2014). The level of fucosylation is of particular significance, since afucosylated IgG has up to 20-fold increased affinity for FcγRIIIa (Dekkers et al., 2017; Ferrara et al., 2011). Even more strikingly, IgG-afucosylation can convert a non-functional ADCC potential to strong and clinically significant responses (Dekkers et al., 2017; Kapur et al., 2014b; Larsen et al., 2021; Shields et al., 2002; Temming et al., 2019; Wang et al., 2017). Increased galactosylation at N297 can further enhance affinity to FcγRIII by additional two fold, and also increases the complement activating capacity of the antibody. In contrast, no influence of bisecting N-acetylglucosamine (GlcNAc) on antibody effector functions has been demonstrated so far (Dekkers et al., 2017).

Fc fucosylation of plasma IgG is near 100% at birth, and although it decreases slightly with age, it normally remains high (∼94%) in adults (Bakovic et al., 2013; de Haan et al., 2016). Nevertheless, very marked and clinically significant reductions (down to ∼10%) in antigen-specific IgG-Fc fucosylation is frequently observed after alloimmunization against erythrocyte and platelet alloantigens (Kapur et al., 2014a; Kapur et al., 2014b; Sonneveld et al., 2017; Sonneveld et al., 2016; Wuhrer et al., 2009). Afucosylation has also been observed for antigen-specific IgG to various enveloped viruses (Ackerman et al., 2013; Larsen et al., 2021; Wang et al., 2017). In human immunodeficiency virus (HIV) infections, low Fc fucosylation has been proposed as a trait of elite controllers (Ackerman et al., 2013), but it is associated with FcγRIIIa mediated immunopathology in SARS-CoV-2- and secondary dengue virus infections (Chakraborty et al., 2021; Larsen et al., 2021; Wang et al., 2017). Vaccination with the attenuated paramyxoviruses measles and mumps also results in specific IgG with reduced fucosylation similar to that acquired after natural infection (Larsen et al., 2021). In contrast, infection with the non-enveloped parvovirus B19, protein subunit vaccination against hepatitis B virus, vaccination with inactivated influenza virus, or vaccination against tetanus, pneumococcal, and meningococcal disease do not induce selectively afucosylated IgG (Larsen et al., 2021; Selman et al., 2012; Vestrheim et al., 2014).

The above findings have led us to propose that afucosylated IgG has evolved as a beneficiary immune response to foreign antigens expressed on host membranes in the context of infections, which is mimicked in alloimmunizations with devastating consequences (Kapur et al., 2014a; Kapur et al., 2015; Kapur et al., 2014b; Larsen et al., 2021; Sonneveld et al., 2017; Sonneveld et al., 2016). In this study, we tested the hypothesis that antibody responses to *P. falciparum* antigens expressed on the IE surface are also a subject to afucosylation. To this end, we examined naturally acquired IgG responses to the PfEMP1 antigens VAR6 and VAR2CSA and to the merozoite antigen GLURP, and VAR2CSA-specific IgG induced by subunit vaccination.

## Results and discussion

### Naturally acquired PfEMP1-specific IgG is highly afucosylated

We first used a set of plasma samples collected from 127 pregnant Ghanaian women at the time of their first visit to antenatal clinics (Ofori et al., 2009), to assess N297 glycosylation of IgG with specificity for three *P. falciparum* recombinant antigens. We used the full ectodomains of VAR2CSA, and the non-pregnancy-restricted Group A-type VAR6, which are both naturally expressed on the IE surface. We also included the merozoite antigen GLURP, which is not expressed on IE surface (Borre et al., 1991) (Fig. 1).

**Figure 1.**
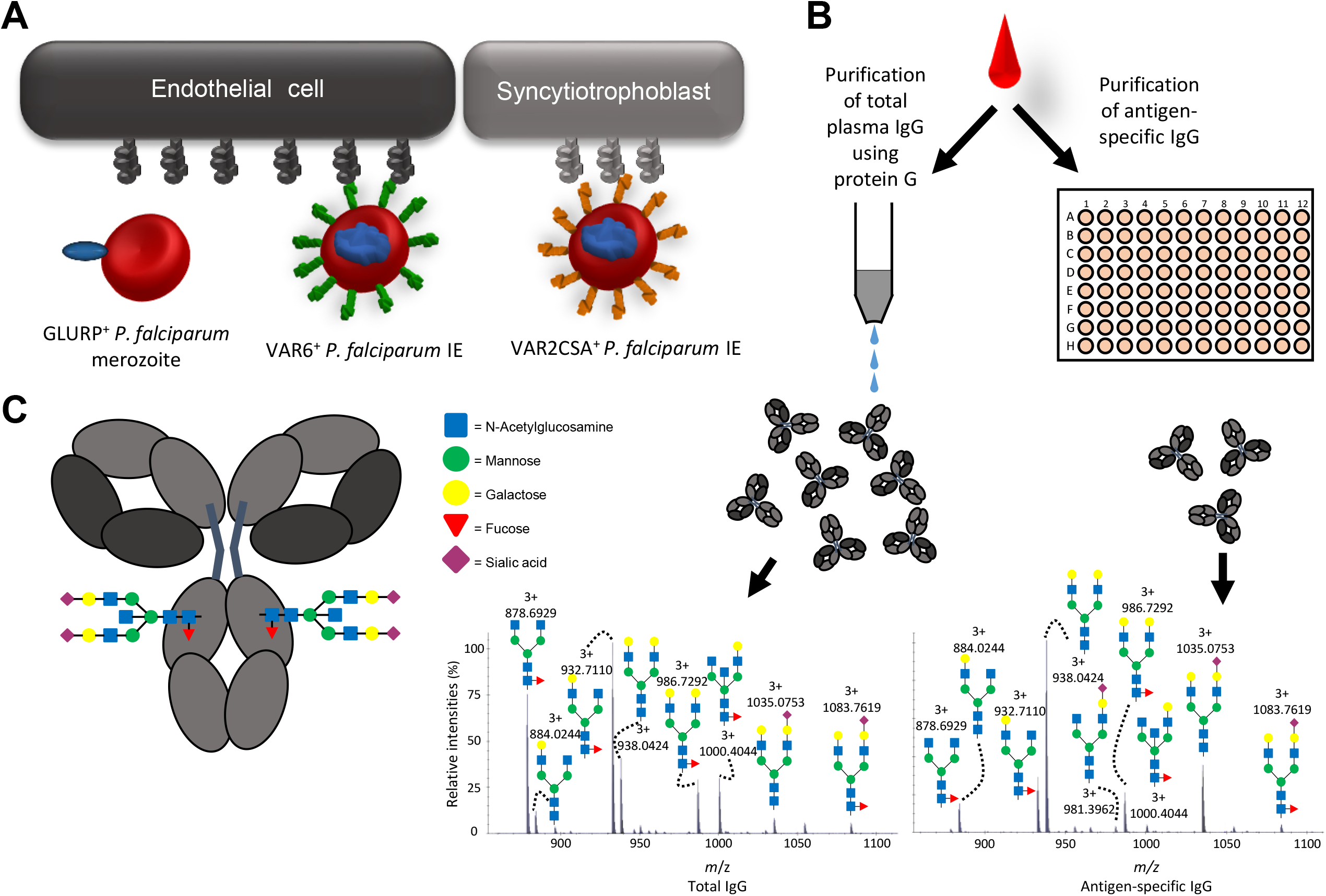
Background and study workflow. **(A)** IgG1 specific for the merozoite antigen GLURP and two members of the PfEMP1 family expressed on the surface of IEs were analyzed in this study. Most PfEMP1 variants facilitate sequestration of IEs to vascular endothelium (exemplified here by VAR6), while VAR2CSA-type PfEMP1 mediate IE sequestration in the placental syncytiotrophoblast and intervillous space. **(B)** Plasma samples were split and used to purify total plasma IgG1 and antigen-specific IgG1, using protein G-coupled sepharose and solid-phase absorption with recombinant antigens, respectively. Eluted IgG1 was digested with trypsin and the glycopeptides analyzed by liquid chromatography mass spectrometry (LC-MS). Examples of MS spectra of total IgG1 (left) and antigen-specific (anti-VAR6) IgG1 (right) from one sample is shown. **(C)** The fractions of the different glycosylation traits of the Fc glycan depicted were calculated from LC-MS spectra.

In line with our hypothesis suggesting that afucosylated IgG response is restricted to foreign antigens expressed on host cells (such as alloantigens and outer-membrane proteins of enveloped viruses (Kapur et al., 2014a; Kapur et al., 2015; Kapur et al., 2014b; Larsen et al., 2021; Sonneveld et al., 2017; Sonneveld et al., 2016)), IgG1-responses to VAR6 and VAR2CSA were markedly Fc afucosylated (Fig. 2A). All individuals showed lowered anti-VAR6 Fc fucosylation compared to total IgG1, which remained high. The magnitude of the decreased Fc fucosylation of VAR6-specific IgG1 exceeded any previously reported pathogen-derived immune response. The most similar responses are against rhesus D on red blood cells and human platelet antigen-1a on platelets.

**Figure 2.**
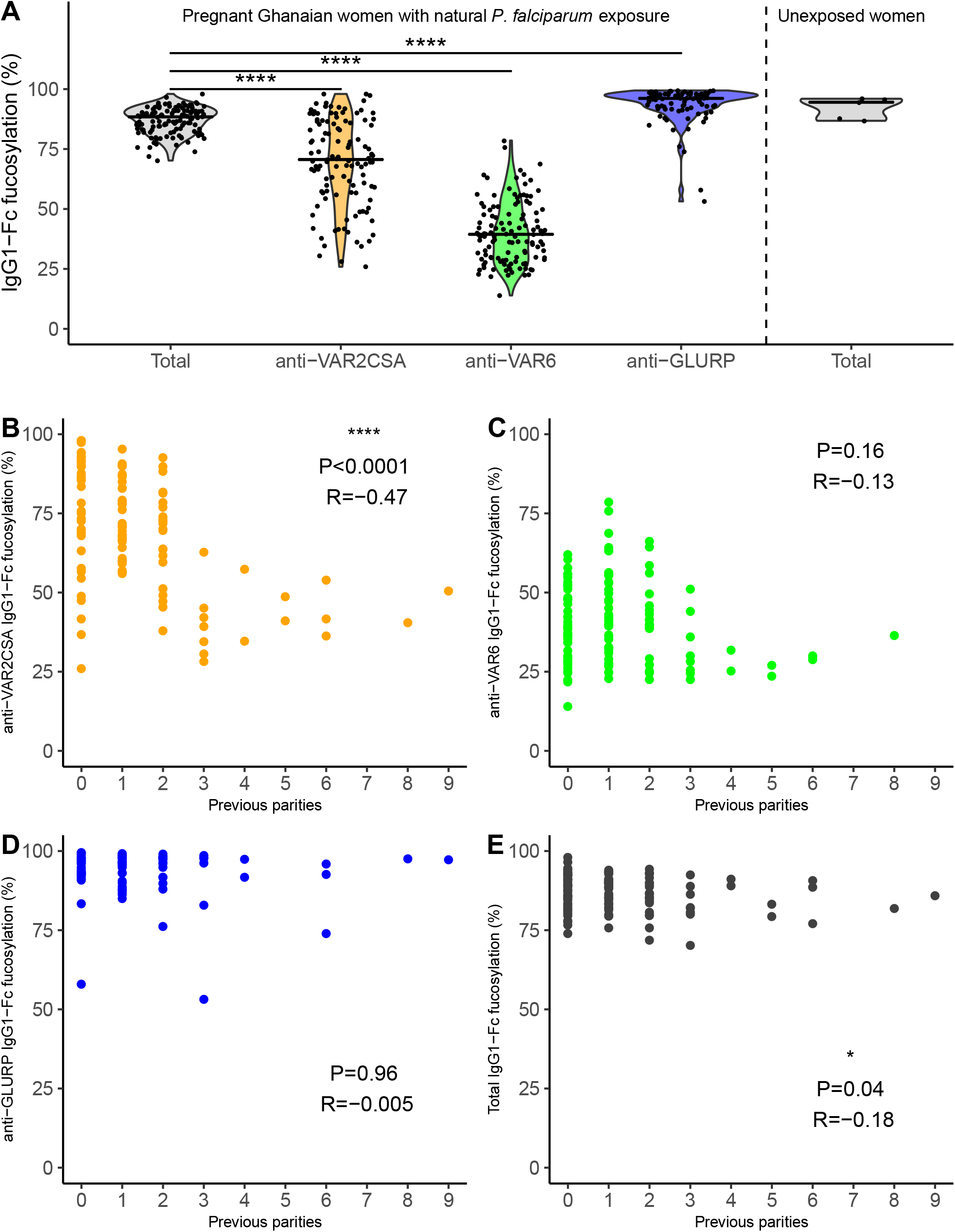
Fc fucosylation of naturally acquired P. falciparum-specific IgG depends on antigen location and exposure. **(A)** Fc fucosylation levels of total plasma IgG1 (gray, n=127) and IgG1 specific for VAR2CSA (orange, n=117), VAR6 (green, n=121), and GLURP (blue, n=88) in Ghanaian pregnant women (left four panels). Fc fucosylation levels of total plasma IgG1 from unexposed Dutch women (n=5) were included for comparison (right panel). Medians and densities are shown. Statistically significant pairwise differences (multiple Wilcoxon signed rank test with Bonferroni correction) are indicated (****: P<0.0001). **(B-E)** Correlations of **(B)** VAR2CSA-, **(C)** VAR6-, **(D)** GLURP-specific and **(E)** total IgG1-Fc fucosylation levels with parity. P-values, and correlation coefficients are shown. Statistical significance of correlations (Spearman’s correlations. *: P<0.05; **: P<0.01; ***: P<0.001; ****: P<0.0001.

However, IgG1 responses to those antigens display big variation in Fc fucosylation ranging from almost 100% to 10% (Kapur et al., 2014a; Kapur et al., 2014b). In contrast, GLURP-specific IgG1 Fc fucosylation was generally high, also in line with our hypothesis (Fig. 2A). A few women showed marked afucosylation of GLURP-specific IgG1 (Fig. 2A), possibly in response to GLURP deposited on the erythrocyte surface during invasion, as has been described for other merozoite-specific antigens (Awah et al., 2009). IgG1 specific for all three *P. falciparum* antigens showed higher Fc galactosylation and sialylation levels than total IgG1, similar to what is known for recent immunizations (Larsen et al., 2021; Selman et al., 2012) (Supplementary Fig. 1A-B). Levels of bisecting GlcNAc were lower for VAR2CSA- and VAR6-specific IgG1, and higher for GLURP-specific IgG1 compared to total IgG1 (Supplementary Fig. 1C). These results indicate that antigen-specific IgG levels are modulated in complex ways according to exposure and antigen context.

Afucosylation of VAR2CSA-specific IgG1 was generally less pronounced than that of VAR6-specific IgG1 (Fig. 2A). Exposure to VAR2CSA-type PfEMP1 occurs later in life, as it is restricted to pregnancy, whereas *P. falciparum* expressing Group A PfEMP1 (such as VAR6) are associated with severe malaria in children (Jensen et al., 2004; Lennartz et al., 2017; Turner et al., 2013). IgG responses to Group A PfEMP1 variants are acquired from early in life in endemic areas through repeated exposure to parasites expressing those variants (Bull et al., 2000; Cham et al., 2009; Cham et al., 2010; Nielsen et al., 2002; Olsen et al., 2018). VAR6-specific IgG1 was consistently afucosylated in all tested individuals, probably as a result of continuous exposure to Group A PfEMP1 in childhood (Fig. 2A), indicating that afucosylation is a persistent phenotype once acquired. In contrast, the level of fucosylation of VAR2CSA-specific IgG1 was more varied (Fig. 2A) and decreased with increased antigen exposure, using parity as proxy (Fig. 2B). This was not the case for VAR6-(Fig. 2C) or GLURP-specific IgG1 (Fig. 2D), and only marginal for total plasma IgG1 (Fig. 2E).

### Fc afucosylation of PfEMP1-specific IgG is stable

The above findings support the hypothesis that afucosylated IgG specific for host membrane-associated immunogens is attained following repeated exposure and that the phenotype is stable once acquired. To examine this hypothesis further, and to consolidate the findings described above, we proceeded to determine the Fc fucosylation of IgG with specificity for the same three antigens, using an availability-based subset (N=72) of plasma samples from a previously published cohort of Ghanaian women sampled while not pregnant (Ampomah et al., 2014a). The findings regarding total and antigen-specific IgG1 (Fig. 3 and Supplementary Fig. 1D-F) were fully consistent with those obtained with the samples from pregnant women. The marked Fc afucosylation of VAR2CSA- and VAR6-specific IgG1 was more pronounced among this second group of women (Fig. 3A), probably reflecting the more intense parasite transmission in the rainforest compared to the coastal savannah where the non-pregnant and pregnant women were recruited, respectively (Ampomah et al., 2014a; Ofori et al., 2009). Although VAR2CSA-specific IgG levels decay markedly within six months of delivery (Ampomah et al., 2014b; Staalsoe et al., 2001), the parity-dependency of the degree of VAR2CSA-specific IgG1 Fc afucosylation remained in these non-pregnant women (Fig. 3B). Furthermore, there was no significant correlation between the time since last pregnancy and Fc fucosylation levels of VAR2CSA-specific IgG1 (Fig. 3C). Taken together, these findings reinforce the inference that PfEMP1-specific IgG1 Fc afucosylation remains stable in the absence of exposure to antigen. This conclusion is in line with our previous findings regarding fucosylation of IgG1 alloantibodies being stable for >10 years (Kapur et al., 2015; Kapur et al., 2014b; Sonneveld et al., 2016). However, unlike the Fc afucosylation of PfEMP1-specific IgG1, which appeared to be exposure-dependent, boosting with alloantigens was found to have no apparent effect on the Fc fucosylation (Kapur et al., 2015; Kapur et al., 2014b; Sonneveld et al., 2016). It also suggests that in these cases, afucosylated IgG1 are secreted by long-lived plasma cells, which for VAR2CSA are sustained for up to a decade after the most recent exposure to parasites expressing VAR2CSA (Ampomah et al., 2014a). This stable response is similar to HIV- and cytomegalovirus-specific responses, but markedly different from initial SARS-CoV-2 responses, which are in most patients only transiently afucosylated for a few weeks after seroconversion (Larsen et al., 2021). This may suggest that those antibodies were either secreted by short-lived plasma cells/plasmablasts, or that afucosylation in those cells is reprogrammed by particular inflammatory conditions.

**Figure 3.**
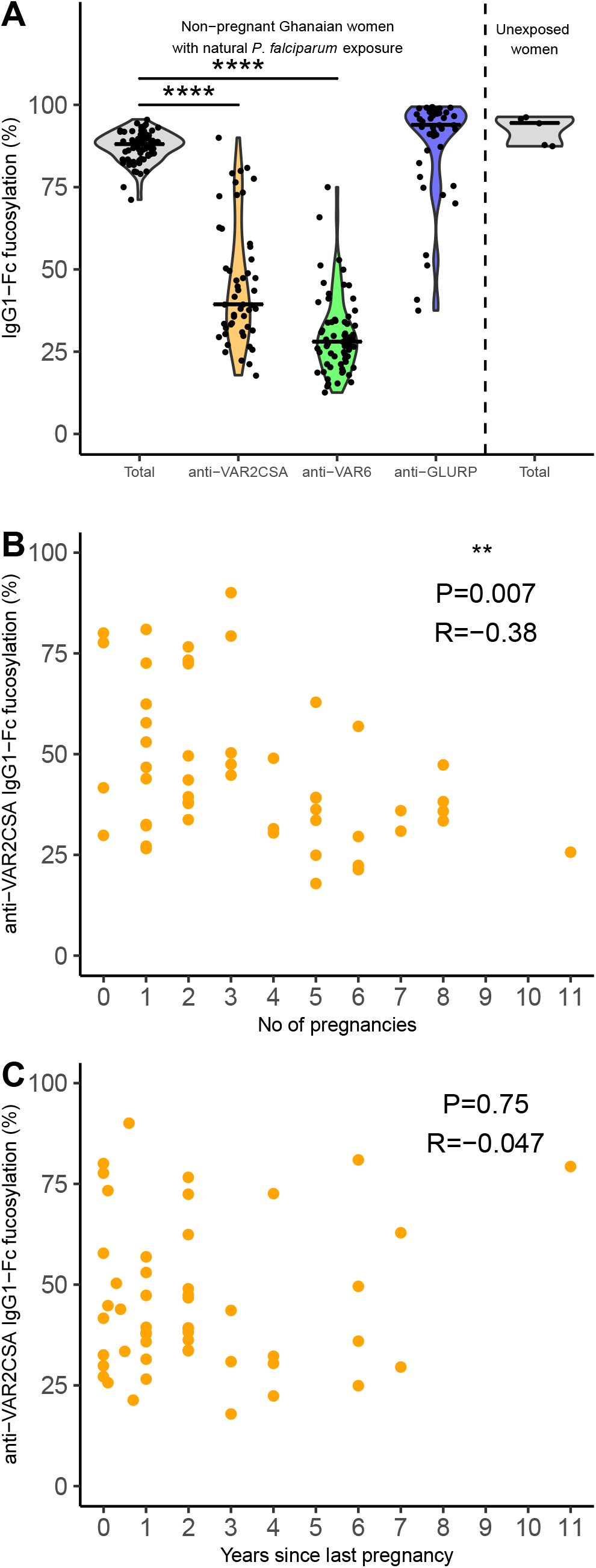
Fc fucosylation levels of VAR2CSA-specifc IgG is temporally stable. Fc fucosylation levels of total plasma IgG1 (gray, n=72) and IgG1 with specificity for VAR2CSA (orange, n=50), VAR6 (green, n=65), and GLURP (blue, n=43) in non-pregnant Ghanaian women exposed to VAR2CSA during one or more previous pregnancies. Fc fucosylation levels of total plasma IgG1 from unexposed Dutch females (n=5) are included as controls. Medians and densities are shown. **(B)** Correlation between fucosylation levels of VAR2CSA-specific IgG1 and parity. **(C)** Correlation between fucosylation levels of VAR2CSA-specific IgG1 and time since last pregnancy. P-values, and correlation coefficients are shown. Statistically significant differences calculated and indicated as in Fig. 2.

### Subunit VAR2CSA vaccination does not induce afucosylated IgG

When measured at the time of delivery, high levels of IgG recognizing placenta-sequestering IEs are strongly associated with protection from adverse pregnancy outcome (Duffy and Fried, 2003; Salanti et al., 2004; Staalsoe et al., 2004). Many of these antibodies interfere with placental IE sequestration (Fried et al., 1998; Ricke et al., 2000), and it is therefore generally assumed that neutralizing (adhesion-blocking) antibodies are required for clinical protection against PM (Beeson et al., 2004; Khunrae et al., 2010; Srivastava et al., 2010). On this basis, development of vaccines to prevent PM, based on the so-called minimal-binding-domain (MBD) of VAR2CSA (Clausen et al., 2012; Srivastava et al., 2011), is currently in progress (Mordmuller et al., 2019; Sirima et al., 2020). To examine the levels of Fc fucosylation of VAR2CSA-specific IgG following subunit vaccination, we tested plasma samples from the PAMVAC Phase 1 clinical trial, which involved adult volunteers without previous *P. falciparum* exposure, vaccinated with a recombinant VAR2CSA-MBD construct (Mordmuller et al., 2019). In contrast to the results obtained with naturally induced VAR2CSA-IgG1, the PAMVAC vaccination induced almost completely fucosylated IgG1, even significantly more fucosylated than total plasma IgG from the same donors (Fig. 4A and Supplementary Fig. 1G-I). This is in line with our recent comparison of naturally acquired and subunit vaccine-induced IgG1 specific for hepatitis B virus (Larsen et al., 2021). To assess the possibility that the full fucosylation of the vaccine-induced VAR2CSA-specific IgG was due to the vaccinees’ lack of previous exposure to *P. falciparum*, genetics, or other environmental parameters, we also tested samples obtained from the parallel trial of the PAMVAC vaccine in Beninese nulligravidae, who were therefore unexposed to VAR2CSA despite lifelong *P. falciparum* exposure. The results (Fig. 4B and Supplementary Fig. 1J-L) were essentially identical to those obtained with unexposed volunteers. Similar to the Ghanaian cohorts described above, the Beninese cohort had lower Fc fucosylation levels of total plasma IgG compared to previous reports of European cohorts and the unexposed vaccine cohort consisting of Europeans, reaffirming previous reports from rural areas (de Jong et al., 2016). This is likely due to accumulating afucosylated IgG to both *P. falciparum* membrane antigens and enveloped viruses (de Haan et al., 2016; Larsen et al., 2021).

**Figure 4.**
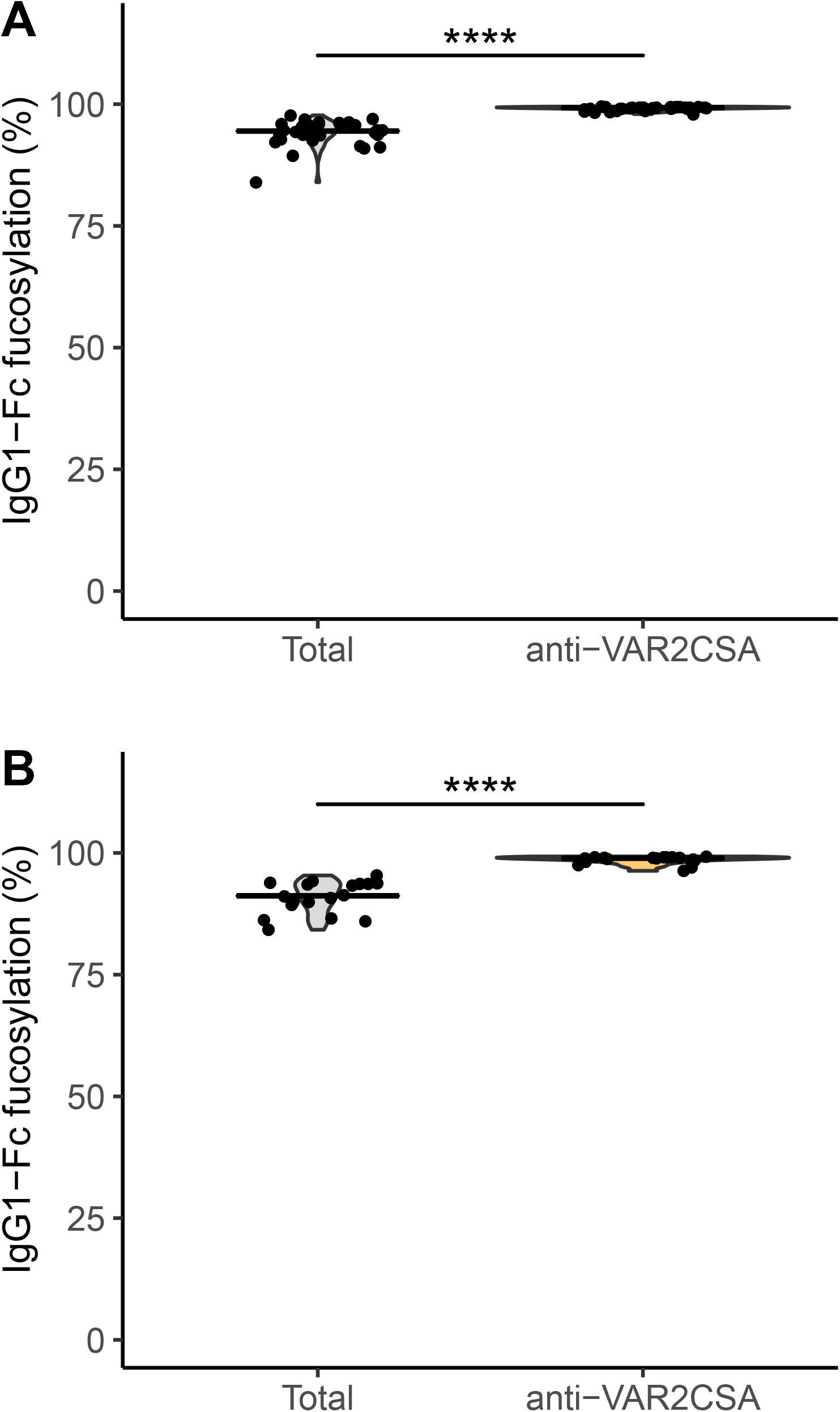
VAR2CSA-specific IgG induced by subunit vaccination is not Fc-afucosylated. Fc fucosylation levels of total (gray) and VAR2CSA-specific (orange) plasma IgG1 in German vaccinees (n=32) without **(A)** and in Beninese vaccinees (n=18) with **(B)** natural exposure to *P. falciparum*. Medians and densities are shown. Statistically significant differences calculated and indicated as in Fig. 2.

### Only afucosylated VAR2CSA-specific IgG induces natural killer cell degranulation

Afucosylation of IgG Fc improves the affinity of IgG for FcγRIII (Dekkers et al., 2017; Ferrara et al., 2011), increasing NK-cell mediated ADCC against IgG-opsonized targets (Temming et al., 2019). Recently it was reported that IgG from individuals naturally exposed to *P. falciparum* makes IEs susceptible to NK-cell mediated ADCC, and that PfEMP1-specific IgG is a major contributor to this response (Arora et al., 2018). To investigate the functional importance of afucosylation of PfEMP1-specific IgG for ADCC, we assessed the ten Ghanaian plasma samples with the highest and lowest Fc fucosylation of VAR2CSA-specific IgG, respectively, for NK cell degranulation efficiency. The samples had a similar distribution of VAR2CSA-specific IgG levels (Fig. 5A). However, they differed markedly in their ability to induce NK-cell ADCC, assessed by degranulation-induced expression of CD107a (Fig. 5B) (Snyder et al., 2018). Only VAR2CSA-specific IgG from individuals with low VAR2CSA-specific Fc fucosylation induced NK-cell degranulation, whereas IgG from individual with high VAR2CSA-specific Fc fucosylation was less effective (Fig. 5B). In line with earlier work (Dekkers et al., 2017; Temming et al., 2019), the fucosylation status of these antibody proved to be a more important predictor of NK-cell mediated activity than their quantity (Fig. 5A). To consolidate these results and to directly compare Fc fucosylation, we next assayed recombinant fucosylation variants of the VAR2CSA-specific human monoclonal antibody PAM1.4. Whereas both bound similarly in ELISA (Fig. 5C), only the afucosylated PAM1.4 induced marked NK-cell degranulation (Fig. 5D). Together, these findings underscore the functional significance of Fc afucosylation of PfEMP1-specific IgG, indicating that IgG induced by PfEMP1 protein subunit vaccination lack potentially important characteristics of the naturally acquired antibody response.

**Figure 5.**
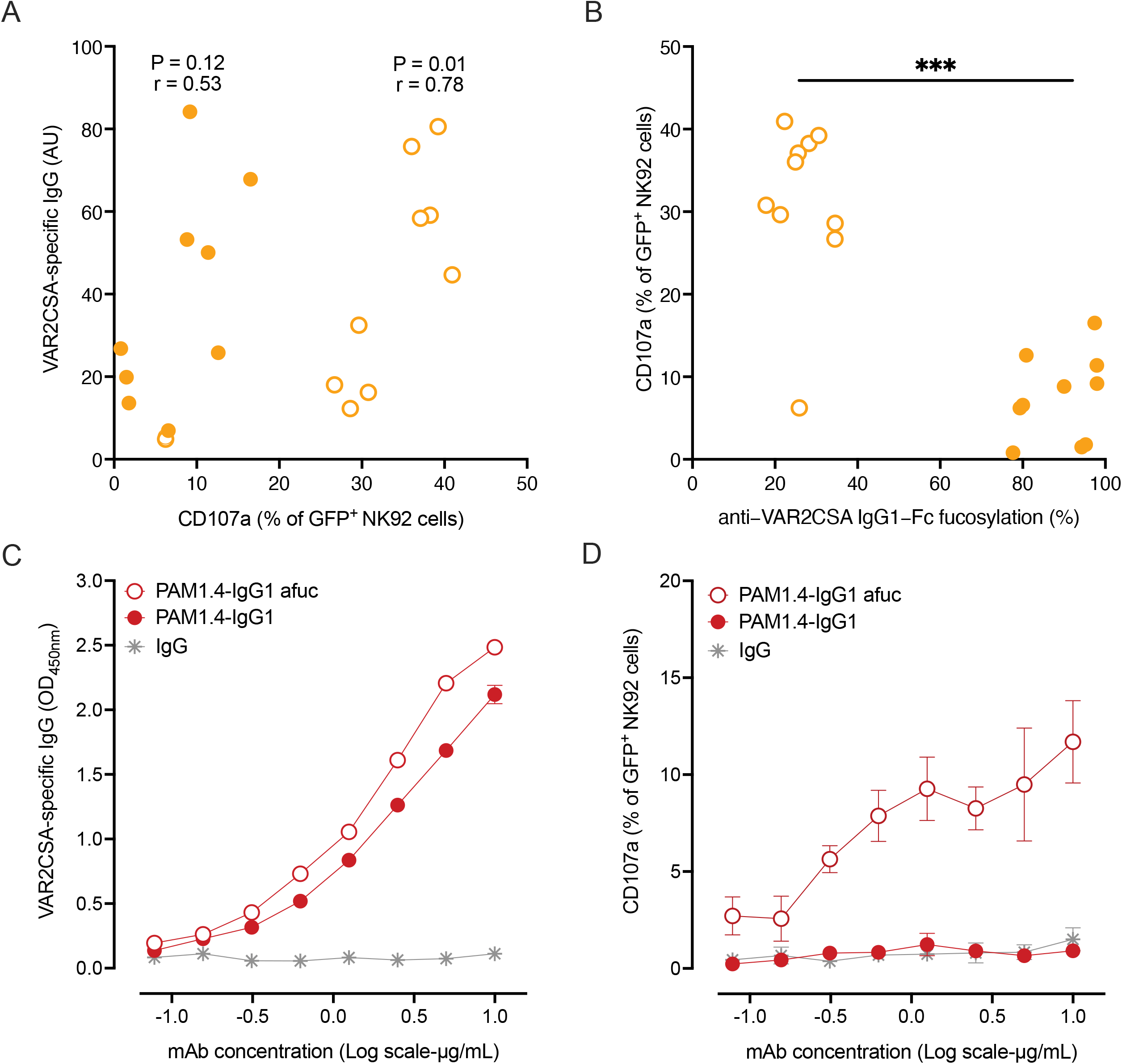
Only afucosylated PfEMP-1 specific IgG induces NK cell-mediated ADCC. Association between **(A)** VAR2CSA-specific IgG levels or **(B)** Fc fucosylation of VAR2CSA-specific IgG and CD107a expression on NK92-CD16a cells. Spearman’s rank correlation (r) and p values are shown for highly fucosylated (filled symbols) and afucosylated anti-VAR2CSA IgG (open symbols) samples, respectively. The groups were compared by Mann-Whitney test. **(C)** Similarly, the VAR2CSA-specific, human monoclonal antibody PAM1.4 as either fucosylated or afucosylated IgG1 was titrated in the same assay and measured for binding or **(D)** degranulation activity (CD107a expression) on NK92-CD16a cells. Data represent mean values ± SD from three independent experiments.

## Conclusion

Our study supports the hypothesis that the immune system has evolved a capacity to selectively modulate the glycosylation pattern of the IgG Fc region, thereby fine-tuning the effector response triggered by antibody-opsonized targets (Larsen et al., 2021). Specifically, it appears that immunogens expressed on host membranes induce afucosylated IgG, thereby increasing its ability to elicit FcγRIII-dependent effector responses such as ADCC. In contrast, immunogens in solution or present on the surface of pathogens seem to mainly induce fucosylated IgG, thus steering the effector response against IgG-opsonized targets towards other FcγR-dependent effector functions. The plasticity in human immune responses to modulate IgG effector functions by altered fucosylation endows the immune system with a so far largely unappreciated level of adaptability.

While it is congruent with the current understanding of how the immune system works, the functional importance of afucosylated IgG in malaria remains to be demonstrated, which future studies will strive to elucidate. In the meantime, it should be emphasized that the decrease in Fc fucosylation reported here exceeds any that has previously been reported for pathogen-derived antigens. Indeed, it also surpasses the clinically significant afucosylation of the IgG response to alloantigens (Kapur et al., 2014a; Kapur et al., 2014b; Wuhrer et al., 2009), thus, implying that the immunopathogenic IgG raised in these instances is an unfortunate mimic of an evolutionary conserved and advantageous immune mechanism against intracellular pathogens. Finally, the data suggest that to induce afucosylated IgG responses with increased ADCC – and potentially protective capacity, alternative vaccination strategies are required, mimicking the expression of antigens on host cells.

## Materials and methods

### Human subjects

We used biological samples collected as part of the following studies: (i) A longitudinal study of malaria in pregnancy, conducted in Dodowa, located in a coastal savannah area with stable, seasonal *P. falciparum* transmission, approximately 40 km North of Accra, Ghana (Ofori et al., 2009). (ii) A cross-sectional study of immune responses to VAR2CSA in healthy non-pregnant women (Ampomah et al., 2014a), conducted in Assin Foso, in a rainforest area with high and perennial *P. falciparum* transmission, located approximately 80 km North of Cape Coast, Ghana (Afari et al., 1995). (iii) A phase 1 clinical trial of the VAR2CSA-based PAMVAC vaccine, conducted in non-immune German volunteers and in adult, nulligravid *P. falciparum*-exposed Beninese women volunteers (Mordmuller et al., 2019). Healthy blood donor samples from Sanquin, Amsterdam, The Netherlands, were used as negative control donors.

The Ghanaian donors all had serologic evidence of exposure to *P. falciparum*, with seropositivity rates above 90% in the non-pregnant cohort (Ampomah et al., 2014a) and above 70% in the pregnant cohort (Data not shown).

A more detailed demographic description of the analyzed cohorts can be found in the supplementary materials (Supplementary table 1).

### P. falciparum *recombinant antigens*

The full-length ectodomains of the VAR2CSA-type PfEMP1 antigen IT4VAR04 (VAR2CSA) and of the Group A PfEMP1 antigen HB3VAR6 (VAR6) were expressed in baculovirus-infected insect cells and purified as described previously (Khunrae et al., 2010; Stevenson et al., 2015). The amino-terminal, non-repetitive R0 region of glutamate-rich protein (GLURP) was expressed in *Escherichia coli* and purified as described elsewhere (Theisen et al., 1995).

### Purification of IgG from plasma samples

Total IgG from individual donors was purified from ∼1 µL plasma using the AssayMAP Bravo platform (Agilent Technologies, Santa Clara, USA) with Protein G-coupled cartridges as described elsewhere (Larsen et al., 2021).

*P. falciparum* antigen-specific IgG was purified from individual donors by incubation (1h, room temperature) of individual plasma samples (diluted 1:10 in phosphate-buffered saline (PBS) supplemented with TWEEN 20 (0.05 %; PBS-T)) in 96-well Maxisorp plates (Nunc, Roskilde, Denmark) coated overnight (4°C; PBS) with VAR2CSA (2 µg/mL), VAR6 (2 µg/mL), or GLURP (1 µg/mL). Following the incubation, the plates were washed 3× with PBS-T, 2× with PBS, and 2× with ammonium bicarbonate (50 mM). Antigen-specific IgG were finally eluted by formic acid (100 mM; 5 min shaking).

### Mass spectrometric IgG Fc glycosylation analysis

Eluates of purified IgG were collected in low-binding PCR plates (Eppendorf, Hamburg, Germany) and dried by vacuum centrifugation (50°C). The dried samples were dissolved in a reduction and alkylation buffer containing sodium deoxycholate (0.4%), tris(2-carboxyethyl)phosphine (10 mM), 2-chloroacetamide (40mM), and TRIS (pH8.5; 100 mM), or ammonium bicarbonate (50 mM). After boiling the samples (10 min; 95°C), trypsin (5 µg/mL) in ammonium bicarbonate (50 mM) was added. The digestion was terminated after overnight incubation (37°C) by acidifying to a final concentration of 2% formic acid. Prior to mass spectrometry injection, sodium deoxycholate precipitates, in samples where this was added, were removed by centrifugation (3,000×g; 30 min), and filtering through 0.65 µm low protein binding filter plates (Millipore, Burlington, USA).

Analysis of IgG Fc glycosylation was performed with nanoLC reverse phase-electrospray-mass spectrometry on an Impact HD quadrupole-time-of-flight mass spectrometer (Bruker Daltonics, Bremen, Germany) and data was processed with Skyline software as described elsewhere (Larsen et al., 2021). The level of fucosylation and bisection were calculated as the sum of the relative intensities of glycoforms containing the respective glycotraits. Galactosylation and sialylation levels were calculated as antenna occupancy. The relative intensities of the glycoforms were summed with mono-galactosylated/sialylated species only contributing with 50 % of their relative intensity.

### Human monoclonal VAR2CSA-specific IgG

The human monoclonal IgG1 antibody, PAM1.4, derived from an EBV-immortalized memory B-cell clone from a Ghanaian woman with natural exposure to PM (Barfod et al., 2007), recognizes a conformational epitope in several VAR2CSA-type PfEMP1 proteins, including IT4VAR04. In the present study, we used a non-modified recombinant version of PAM1.4 produced in HEK293F cells with high Fc fucosylation and a glyco-engineered variant with low Fc fucosylation (Dekkers et al., 2016; Larsen et al., 2019).

### Quantification of VAR2CSA-specific IgG

Levels of VAR2CSA-specific IgG were assessed by ELISA as previously described (Lopez-Perez et al., 2018). In brief, 96-well flat-bottom microtiter plates (Nunc MaxiSorp, Thermo Fisher Scientific) were coated overnight at 4°C with full-length VAR2CSA (100 ng/well in PBS. Monoclonal antibody (0.08 to 10 μg/mL) or plasma samples (1:400) were added in duplicate, followed by washing and horseradish peroxidase-conjugated rabbit anti-human IgG (1:3,000; Dako). Bound antibodies were detected by adding TMB PLUS2 (Eco-Tek), and the reaction stopped by the addition of 0.2 M H_2_SO_4_. The optical density (OD) was read at 450 nm and the specific antibody levels were calculated in arbitrary units (AU), using the equation 100 × [(OD_SAMPLE_-OD_BLANK_)/(OD_POS.CTRL_-OD_BLANK_)].

### Antibody□dependent cellular cytotoxicity (ADCC) assay

Degranulation-induced CD107a expression in response to IgG bound to plastic-immobilized antigen is a convenient marker of NK-cell ADCC (Jegaskanda et al., 2013). Here, we coated 96-well flat-bottom microtiter plates (Nunc MaxiSorp; Thermo Fisher Scientific) overnight at 4°C with full-length VAR2CSA (100 ng/well in PBS; (Lopez-Perez et al., 2018)).Following 1h blocking with PBS containing 1% Ig-free bovine serum albumin-BSA (1% PBS-BSA), plasma samples (1:20) or PAM1.4 variants (0.08 to 10 µg/mL) were added for 1h at 37°C. After washing, 1.6×10^5^ NK92 cells stably expressing CD16a and GFP (Snyder et al., 2018) were added to each well. In addition, anti-human CD107a-PE (H4A3 clone; BD Biosciences), 10 μg/mL brefeldin A (Sigma-Aldrich), and 2 µM monensin (Sigma-Aldrich) were added, and the cells incubated for 4 h at 37°C. Cells were then centrifuged and stained with near-IR fixable Live/Dead dye (Invitrogen), followed by data acquisition on a FACS LSRII flow cytometer (BD Biosciences), and analysis with FlowLogic software (Inivai Technologies, Australia). Wells with antigen and NK cells, but without antibody were included in all experiments to control for unspecific activation. Plasma samples from four Danish non-pregnant women without malaria exposure and purified human IgG (Sigma-Aldrich) were included as negative controls.

### Statistical tests

Statistical analyses were performed using R: A Language and Environment for Statistical Computing (Version 3.5.2). Performed tests are mentioned in the text.

### Ethics statement

Collection of biological samples for this study was approved by the Institutional Review Board of Noguchi Memorial Institute for Medical Research, University of Ghana (study 038/10-11), by the Regional Research Ethics Committees, Capital Region of Denmark (protocol H-4-2013-083), by the Academic Medical Center Institutional Medical Ethics Committee of the University of Amsterdam, by the Ethics Committee of the Medical Faculty and the University Clinics of the University of Tubingen, and by the German Regulatory authorities. The study was conducted in adherence with the International Council for Technical Requirements for Human Use guidelines and the principles of the Declaration of Helsinki. Written informed consent was obtained from all participants before enrollment.

## Author contributions

<u>Conceptualization</u>: MDL, CEvdS, LH, GV

<u>Funding acquisition</u>: MFO, LH, GV

<u>Investigation</u>: MDL, MLP, JN, MW, LH, GV

<u>Study materials</u>: EKD, PA, BM, AS, MAN, MFO, NTN, AM

<u>Writing – original draft</u>: MDL, LH, GV

<u>Writing – review & editing</u>: All.

## Acknowledgments

We are grateful to all the individuals donating blood samples for this study, and to the scientists and health workers participating in the studies for which they were originally collected. We thank Michael Theisen (University of Copenhagen and Statens Seruminstitut) for GLURP antigen and GLURP-reactive IgG preparation, and Bruce Walcheck and Geoff Hart (University of Minnesota) for the NK92-CD16a cell line. We also acknowledge Erik de Graaf (Sanquin Research) for optimization of the analysis pipelines used in this study, done in relation to previous projects.

The study was funded by the Landsteiner Foundation for Blood Transfusion Research grant number 1721 and the Danish International Development Agency (Danida), 12-081RH and 17-02-KU). The PAMVAC study (ClinicalTrials.gov ID NCT02647489) was sponsored by the Universitätsklinikum Tübingen and funded by the European Union Seventh Framework Programme (FP7-HEALTH-2012-INNOVATION; under grant agreement 304815), the Danish Advanced Technology Foundation (under grant number 005-2011-1), and a Medium Scale Collaborative Project supported by the German Federal Ministry of Education and Research (Bundesministerium für Bildung und Forschung) through EVI, KfW, and Irish Aid. The funders had no role in study design, data collection and analysis, decision to publish, or preparation of the manuscript.

## Abbreviations

ADCC: Antibody-dependent cellular cytotoxicity
CM: Cerebral malaria
Fc: fragment crystallizable
FcγR: Fcγ receptor;
GlcNac: N-acetylsglucosamin
GLURP: Glutamate-rich protein
IgG: Immunoglobulin G
IE: Infected erythrocyte
MBD: Minimal-binding domain
PBS: Phosphate-buffered saline
PBS-T: PBS supplemented with TWEEN20
PfEMP1: *Plasmodium falciparum* erythrocyte membrane protein-1
PM: placental malaria.

---

**Supplementary Figure 1. Fc glycosylation traits of P. falciparum-specific IgG in pregnant women**

**(A, D, G, and J)** Fc galactosylation-, **(B, E, H, and K)** Fc sialylation-, and **(C, F, I, and J)** Fc bisecting GlcNAc levels of total IgG1 (gray) and IgG1 with specificity for VAR2CSA (orange), VAR6 (green), and GLURP (blue) in **(A to C)** pregnant Ghanaian women, **(D to F)** non-pregnant Ghanaian women, **(G to I)** *P. falciparum*-naïve German and **(J to L)** VAR2CSA-naïve Beninese vaccinees. Medians and densities are shown. Statistically significant differences calculated and indicated as in Fig. 2.

**Supplementary Table 1.** Summary statistics of plasma donors studies.

Summary statistics of plasma donors studies
| Cohort | Origin | Donors (N) | Women (N; %) | Age (median; inter-quartile range (in years)) | Ref. |
| --- | --- | --- | --- | --- | --- |
| *P. falciparum*-exposed and pregnant | Ghana | 127 | 127; 100% | 24; 20-27 | (Ofori et al., 2009) |
| *P. falciparum*-exposed and non-pregnant women | Ghana | 72 | 72; 100% | 29; 23-38 | (Ampomah et al., 2014a) |
| Non-exposed vaccinees | Germany | 36 | n.a.<sup>1</sup> | Adults<sup>1</sup> | (Mordmuller et al., 2019) |
| *P. falciparum*-exposed vaccinees | Benin | 21 | 21; 100% | Adults<sup>1</sup> | (unpublished) |

<sup>1</sup> Data not available due to blinding of the clinical trial data

**Supplementary Table 2.** Overview of included Fc glycopeptides.

Overview of included Fc glycopeptides
| N-Glycopeptide | m/z 2+ | m/z 3+ | Retention time (sec) |
| --- | --- | --- | --- |
| IgG1 H3N4F1S0 [G0F] | 1317.526 | 878.687 | 80 |
| IgG1 H4N4F1S0 [G1F] | 1398.552 | 932.704 | 78 |
| IgG1 H5N4F1S0 [G2F] | 1479.579 | 986.722 | 77 |
| IgG1 H3N5F1S0 [G0FN] | 1419.066 | 946.380 | 81 |
| IgG1 H4N5F1S0 [G1FN] | 1500.092 | 1000.398 | 79 |
| IgG1 H5N5F1S0 [G2FN] | 1581.119 | 1054.415 | 78 |
| IgG1 H3N4F0S0 [G0] | 1244.497 | 830.001 | 83 |
| IgG1 H4N4F0S0 [G1] | 1325.524 | 884.018 | 82 |
| IgG1 H5N4F0S0 [G2] | 1406.550 | 938.036 | 81 |
| IgG1 H3N5F0S0 [G0N] | 1346.037 | 897.694 | 83 |
| IgG1 H4N5F0S0 [G1N] | 1427.063 | 951.712 | 82 |
| IgG1 H5N5F0S0 [G2N] | 1508.090 | 1005.729 | 79 |
| IgG1 H4N4F1S1 [G1FS] | 1544.100 | 1029.736 | 77 |
| IgG1 H5N4F1S1 [G2FS] | 1625.127 | 1083.754 | 75 |
| IgG1 H4N5F1S1 [G1FNS] | 1645.640 | 1097.429 | 77 |
| IgG1 H5N5F1S1 [G2FNS] | 1726.667 | 1151.447 | 77 |
| IgG1 H4N4F0S1 [G1S] | 1471.071 | 981.050 | 80 |
| IgG1 H5N4F0S1 [G2S] | 1552.098 | 1035.068 | 79 |
| IgG1 H4N5F0S1 [G1NS] | 1572.611 | 1048.743 | 77 |
| IgG1 H5N5F0S1 [G2NS] | 1653.638 | 1102.7610 | 77 |
| IgG1 H5N4F1S2 [G2FS2] | 1770.675 | 1180.786 | 76 |

