## Supplementary figures and images for "Afucosylated *Plasmodium falciparum*-specific IgG is induced by infection but not by subunit vaccination"

### Supplemental figure 1

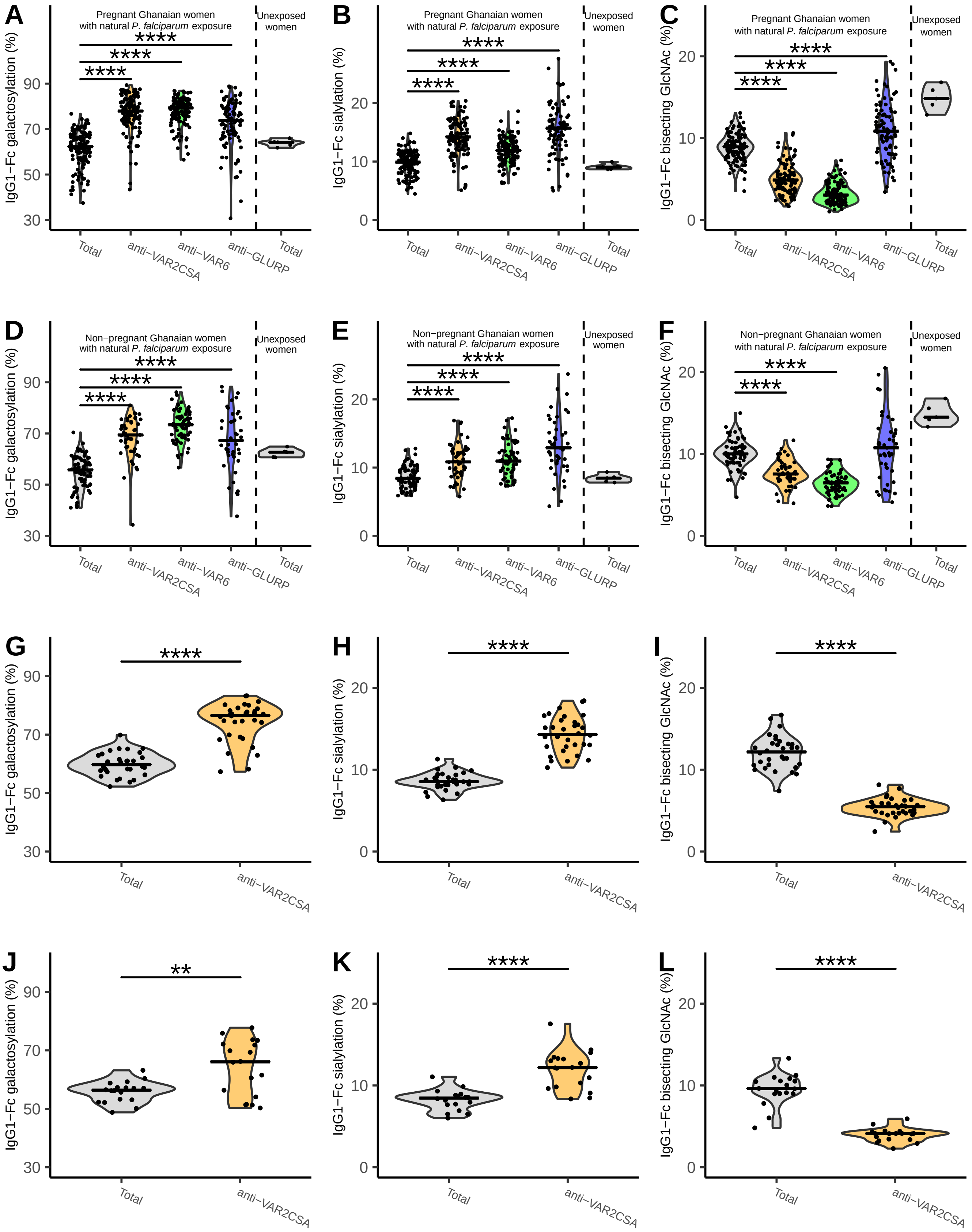
